# *Drosophila* Sociality Influences Immune Peptide Load and Apoptosis-induced Tumor Suppression independently of the Antitumor Peptide Defensin

**DOI:** 10.1101/2025.10.21.683679

**Authors:** Pierre Delamotte, Perla Akiki, Mickaël Poidevin, Delphine Naquin, Maxence de Taffin de Tilques, Xingyi Cheng, Frederic Mery, Frederic Marion-Poll, Jacques Montagne

## Abstract

Patient’s social environment might influence cancer outcome. This potentially happens in *Drosophila*, as we previously reported that the social context influences the growth of intestinal tumors. To uncover the underlying social-induced physiological mechanisms, we performed RNA-seq of isogenized beheaded tumorous *Drosophila*. Importantly, expression of several immune peptides varied according to the social-induced tumor growth effect. Furthermore, ectopic expression in tumors of the apoptotic-inhibitor p35 suppressed the social-induced effect. Next, we challenged the immune peptide Defensin, previously reported to suppress imaginal disc tumor growth through a cell-death/JNK-dependent network. Nonetheless, the social-induced tumor suppression was maintained upon Defensin overexpression or JNKK-knockdown in tumors, and in *defensin* mutants. Surprisingly, tumor growth was reduced in the latter, indicating that Defensin sustains the growth of these intestinal tumors. In summary, our study indicates that the social context affects the immune response and that a given immune peptide may have opposite effects depending on tumor type.

## INTRODUCTION

Studies in humans to evaluate the impact of sociality and wellbeing on health and disease suffer from the complexity of intercorrelated factors, leading to potential uncontrolled bias (Mcclintock *et al*. 2005; Chida *et al*. 2008; Hermes *et al*. 2009; Cenat *et al*. 2023; Fontesse *et al*. 2023). Animal models provide alternative powerful systems to address these issues. Using the *Drosophila* model, we previously reported a social incidence on tumor growth in females bearing genetically induced intestinal tumors (Dawson *et al*. 2018).

Social-induced tumor-suppression, if effective in humans, might act through various physiological processes. Social stresses have been reported to impact endocrine- and metabolic-based diseases in rodents (Nonogaki *et al*. 2007; Lin *et al*. 2015), while metabolic homeostasis is tightly connected to cancer progression (Altea-Manzano *et al*. 2025). The growth hormone/insulin-like growth factor-1 (GH/IGF1) axis-dependent regulation of growth and homeostasis may have a direct effect on tumor progression (Werner and Laron 2023). Further, cancerous patients commonly experience death anxiety and depression, justifying the need for psychological support (Brown *et al*. 2025; Tang *et al*. 2025). Imbalanced immunity has been associated with depression (Matsuo *et al*. 2025), while the immune system is known to impact cancer progression (Marcus *et al*. 2014; Sharma and Allison 2015a; Iwai *et al*. 2017; Olawade *et al*. 2025). These studies indicate that a complex network of physiological functions may control cancer progression, involving patient health and interactions with the tumor microenvironment (Anderson and Simon 2020).

Despite the rare occurrence of spontaneous tumors in testis and intestine (Salomon and Jackson 2008), several experimental tumor models have been developed in *Drosophila* (Brumby and Richardson 2005; Montes-Labrador *et al*. 2025). Clonal expression in imaginal discs ─larval anlage of adult structures─ of the RasV12 oncogene combined with *scribble* loss-of-function results in metastatic tumors (Pagliarini and Xu 2003). Ras/Src tumors in imaginal discs become aggressive in larvae fed high sugar diet, while most tissues display insulin resistance (Hirabayashi *et al*. 2013). Eiger, the unique *Drosophila* member of the Tumor-necrosis-factor superfamily, which acts via JNK signaling (Igaki *et al*. 2002), may have opposite effects depending on the model (Moreno *et al*. 2002; Brumby and Richardson 2003; Uhlirova *et al*. 2005; Igaki *et al*. 2006; Uhlirova and Bohmann 2006; Igaki *et al*. 2009). The *Drosophila* immune system relies on hemocytes and immune peptides mainly produced in the fat body, an insect organ that fulfills adipose, hepatic and immune functions (Lemaitre and Hoffmann 2007; Moraes and Montagne 2021). It has been shown that transformed imaginal discs respond to an Eiger-induced tumor suppression involving hemocytes and fat body (Parisi *et al*. 2014). This results from the recruitment of the immune peptide Defensin that triggers apoptosis to tumor cells (Parvy *et al*. 2019), thereby proving the fruitfly a powerful model to investigate systemic regulation of cancer growth.

We previously reported that the growth of intestinal tumors was reduced for flies maintained in a homogenous group compared to flies maintained alone either in isolation or with control healthy flies (Dawson *et al*. 2018). RNAseq from the head of these tumorous flies revealed social-dependent changes in gene expression related to nervous system activity, suggesting that *Drosophila* have a cognitive perception of their social environment (Akiki *et al*. 2024). In our present study, RNAseq from the body of tumorous flies reveals that the social context provokes changes in the expression of genes encoding immune peptides, suggesting that the immune system is responsible for the social-induced tumor suppression. While previous studies reported the antitumoral effect of JNK signaling and immune peptide Defensin (Parisi *et al*. 2014; Parvy *et al*. 2019), we observed that the tumors suppression depends on apoptosis but neither on JNK signalling nor on Defensin. Unexpectedly, we observed that in *defensin* mutant flies tumor growth is overall reduced, suggesting that Defensin has a protumoral impact on our intestinal tumor model.

## METHODS & MATERIALS

### Fly stocks and husbandry

The *yw,HS-flp*;*esg-gal4,UAS-GFP*; *FRT82B,Tub-Gal80* (line 1) and *w,HS-flp*;*UAS-Ras*^*V12*^;*FRT82B,Apc2*^*N175K*^,*Apc*^*Q8*^ (line 2) stocks (Martorell *et al*. 2014) were maintained with co-segregating *SM5-TM6B* balancers. Prior to experiments, these lines were isogenized on chromosomes I, II and III from individual males crossed to a *FM7; SM5-TM6B* stock. Next, the *UAS-Ras*^*V12*^ chromosome II from line 2 was recombined with either *UAS-p35* (BDSC#5072), *UAS-JNKK-RNAi* (VDRC#109277), *UAS-Defensin* or the *def*^*sk3*^ mutant allele (Parvy *et al*. 2019), the latter being also recombined with the *esg-gal4,UAS-GFP* chromosome II from line 1; these recombined chromosomes were re-introduced into the parental lines. Control *w*^*1118*^ flies were used as healthy females in H-groups.

To generate tumors, females from line 1 or its derivative recombined to the *def*^*sk3*^ mutation were crossed to males from line 2 or its recombined derivatives. In their progeny non-balanced virgin females were selected and heat-shocked at 37°C for 1h15mins, two to three days after adult eclosion as previously detailed (Delamotte *et al*. 2025). Fly groups were established immediately after heat-shock and maintained in a 25°C incubator on our standard media (Garrido *et al*. 2015), changed twice a week into new vials.

### Midgut dissection and flow cytometry

24 days past heat-shock, midguts from individual flies were dissected and dissociated into single cells as previously described (Akiki *et al*. 2024). Samples containing the intestinal cells of single (Fig. 4 and Table S6) or group of two to three (Fig. 5 and Table S7) flies were processed on a CytoFLEX S (Beckman Coulter) cytometer and analyzed with CytExpert 2.4. Results are presented as mean percentages of tumorous cells (labelled by GFP) over the total intestinal cell number (inclusing tumorous cells). Statistics were conducted on GraphPad Prism 8.0.1. using a Mann-Whitney test.

### Transcriptome analyses (RNAseq)

For each replicate, groups of 100 tumorous flies were beheaded on dry ice; heads and bodies were store separately at -80°C. These flies were selected 10 days post tumor induction, as we assumed that the social-induced effect occur along the entire process of tumor growth. RNA extraction from bodies was performed as described for heads (Akiki *et al*. 2024). Briefly, RNAs were extracted using Trizol-chloroform and libraries for Illumina sequencing were prepared using the Illumina TruSeq total RNA stranded kit (Garrido *et al*. 2015) and sequenced using a NextSeq 550 instrument. DNA reads were converted into raw data (FASTQ files) and preprocessed to remove adapters, low-quality sequences and artefacts. Cleaned sequences were aligned to the reference genome “dmel-all-chromosome-r6.42.fasta” using HISAT2. Differential analyses between replicates were performed using the DESeq2 tool to identify genes whose expression significantly differs between experimental conditions. Output of DeSeq2 was finally analyzed with in-house Python scripts. The gene set enrichment analysis was performed using FlyEnrichr (Chen *et al*. 2013; Kuleshov *et al*. 2016).

## RESULTS

### RNA-Seq of tumoral body revealed social-induced variations

In our early study (Dawson *et al*. 2018), we reported that the growth of genetically-induced intestinal tumors was reduced when eight tumorous *Drosophila* virgin females were maintained in a homogenous group (G-flies), compared to tumorous females maintained alone, either in isolation (A-flies) or in a heterogeneous group with seven non-tumorous control females (H-flies) (Dawson *et al*. 2018). More recently, we performed head transcriptome analysis from tumorous flies (three replicates per social condition, each containing 100 heads) collected 10 days after tumor induction, since tumor growth significantly increases between days 7 and 14 post-tumor induction (Akiki *et al*. 2024). In the present study, the corresponding bodies were used for transcriptome analysis to identify the putative physiological processes induced by the social context, which could modulate tumor growth in a systemic manner. Given the isogenic nature of the fly lines, differences between conditions results solely from the social context in which flies are distributed after tumor induction. Congruent to the RNAseq from fly heads (Akiki *et al*. 2024), principal component analysis revealed distinct segregation of the three replicates of H(heterogeneous)-flies from the three replicates of G(group)-flies, whereas A(alone)-replicates segregate in between the G- and H-replicates (Fig. 1A). PC1 was the key parameter to separate social-dependent differences, while PC2 revealed potential variability among H-replicates. Differentially expressed genes were selected based on a P_adjusted_ <0.05 and a Log2foldchange >│1│. Compared to G-replicates, out of 9637 genes evaluated, 248 were found to be significantly up-regulated and 169 significantly down-regulated in H-replicates (Table S1 and S2, and Fig. S1). In A-replicates, compared to G-replicates, 50 genes were found to be significantly up-regulated and 44 genes significantly down-regulated (Table S3 and S4, and Fig. S1). In contrast, no significant differences in gene expression were observed when comparing H-to A-flies (Table S5). Volcano plots visually represent these significant differences in gene expression when comparing H- or A-to G-flies, while no differences are evident when comparing H-to A-flies (Fig. 1B-D).

**Figure 1:**
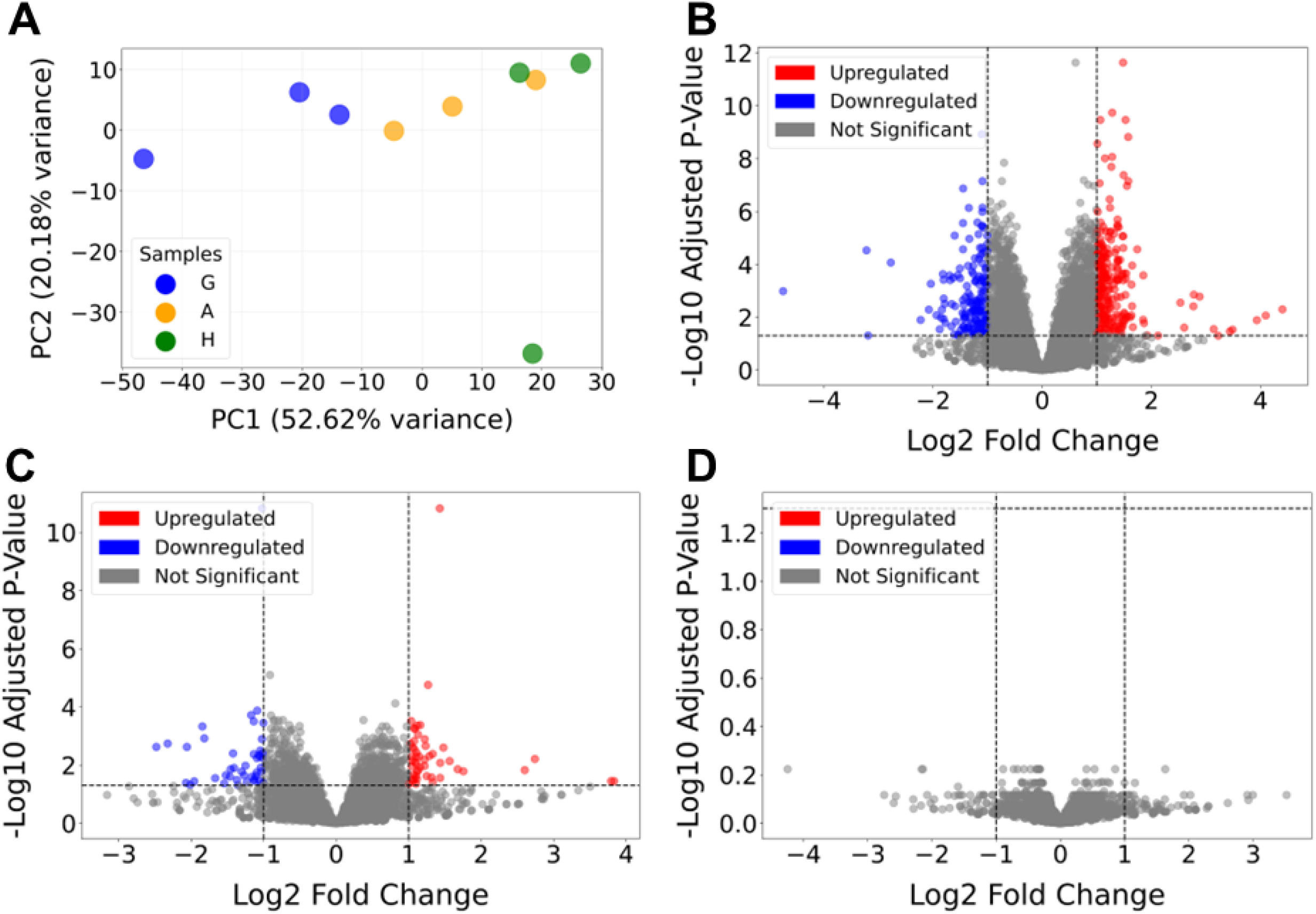
Transcriptomes (RNAseq) from the bodies of G-H- and A-tumorous flies. (**A**) Principal components analysis comparing G(blue)-, H(green)- and A(yellow)-replicates, representing <u>H</u>eterogeneous, <u>A</u>lone or homogenous <u>G</u>roup, respectively. (**B-D**) Volcano plots displaying significantly upregulated (red) and downregulated (blue) genes when comparing H-to G-conditions (**B**), A-to G-conditions (**C**) and H-to A-conditions (**D**).

Gene ontology analysis of the gene candidates using FlyEnrichR revealed that the functions of genes upregulated in H- or A-*versus* G-flies were related to nervous system activity, behavior and cell adhesion (Fig. 2A-B). Additionally, the functions of genes downregulated in H- or A-*versus* G-flies were related to several different biological processes, including early development, morphogenesis and cellular activity (Fig. 2C,D). Importantly, the antibacterial response was found downregulated in both A- and H-flies (Fig. 2C,D).

**Figure 2:**
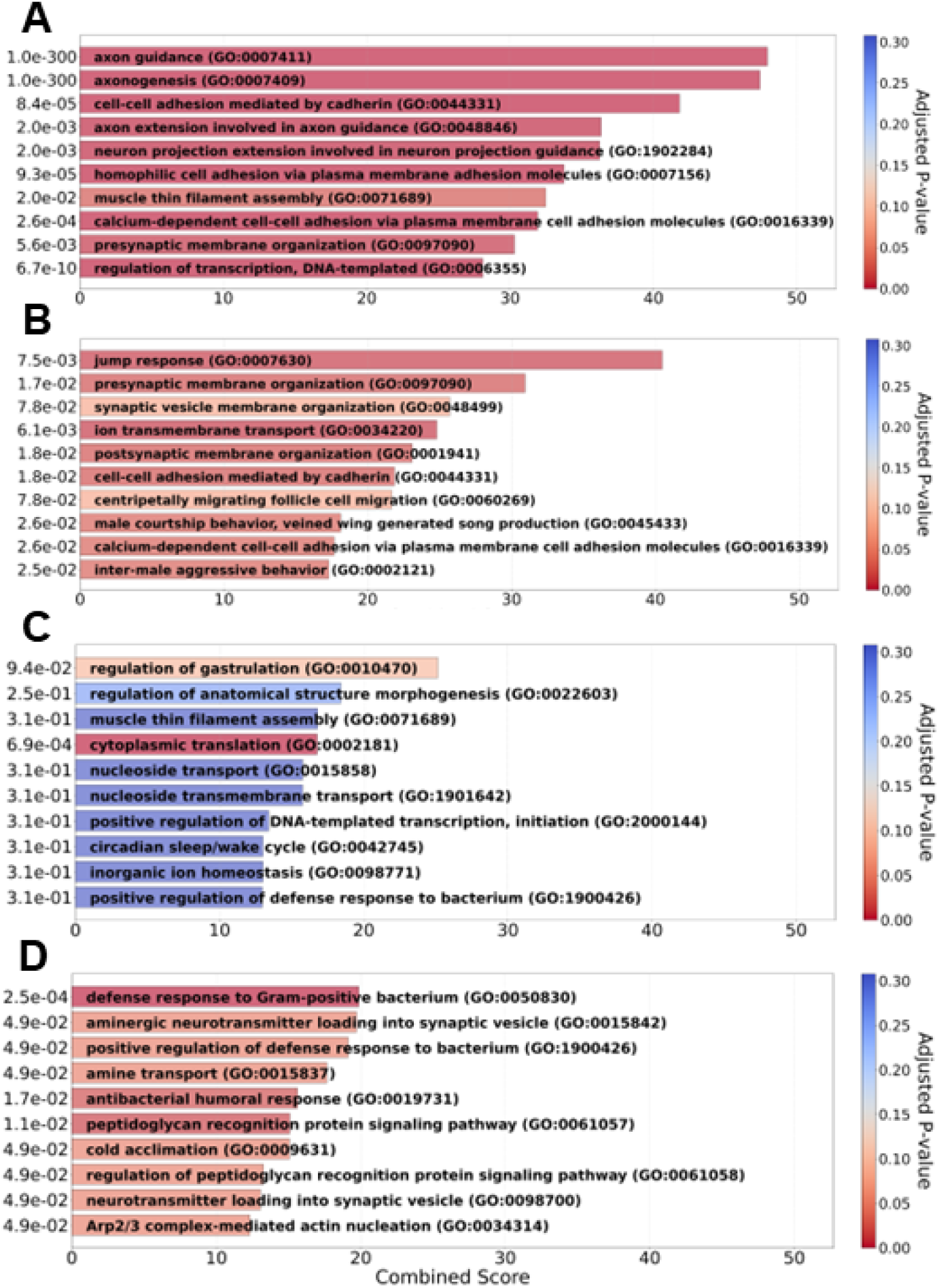
Transcriptome analysis of genes up- or down-regulated in the bodies of H- and A-compared to G-tumorous flies. Biological processes identified while analyzing the genes significantly upregulated (**A**,**B**) and downregulated (**C**-**D**) in the bodies of H(**A**,**C**)- and A(**B**,**D**)-compared to G-flies.

Next, we focused on various processes potentially involved in systemic regulation of growth, including i) energy metabolism (anaerobic glycolysis, TCA and OXPHOS), ii) stress response, iii) immune peptides, and iv) Imd, toll, JNK and insulin signaling pathways (Fig 3). In this way, a tendency could potentially be observed in volcano plots with lower P-value supporting a particular involvement of the challenged biological function. Consistently, antimicrobial peptides (AMPs) focused analysis revealed significant downregulation in A- and H-compared to G-flies (Fig. 3A). The Imd and Toll pathways play a central role in the transcriptional control of genes involved in the *Drosophila* immune response (De GREGORIO *et al*. 2002). However, the expression of intermediate genes of these two pathways was not significantly changed regarding the social context (Fig. 3B-C), suggesting that subtle changes in the activity of these two pathways are sufficient to affect immune peptide gene expression. Alternatively, we cannot exclude that the differences in immune peptide gene expression may rely from a yet unidentified regulation independent of the Imd or Toll pathways. Furthermore, no significant change in gene expression could be observed when analyzing the JNK and insulin signaling pathways and the response to oxidative stress, although genes of energy metabolism tended, but not significantly, to be downregulated in A- and H-flies (Fig. 3D-G). Taken together, these findings exclude a potential role of insulin-like related growth or oxidative stress, but rather suggest that the immune response might be responsible for the social-dependent tumor-suppression. Nonetheless, it is questionable whether the non-significant changes in the expression of genes acting on energy metabolic homeostasis might be responsible for the social-dependent effect. Finally, considering the critical epithelial-mesenchymal transition step in cancer progression (Beltzner and Pollard 2004; Dongre and Weinberg 2019) and its requirement to sustain the growth of this tumor model (Beltzner and Pollard 2004; Delamotte *et al*. 2025), we cannot neglect a potential role of these genes in mediating the social-dependent tumor suppression.

**Figure 3:**
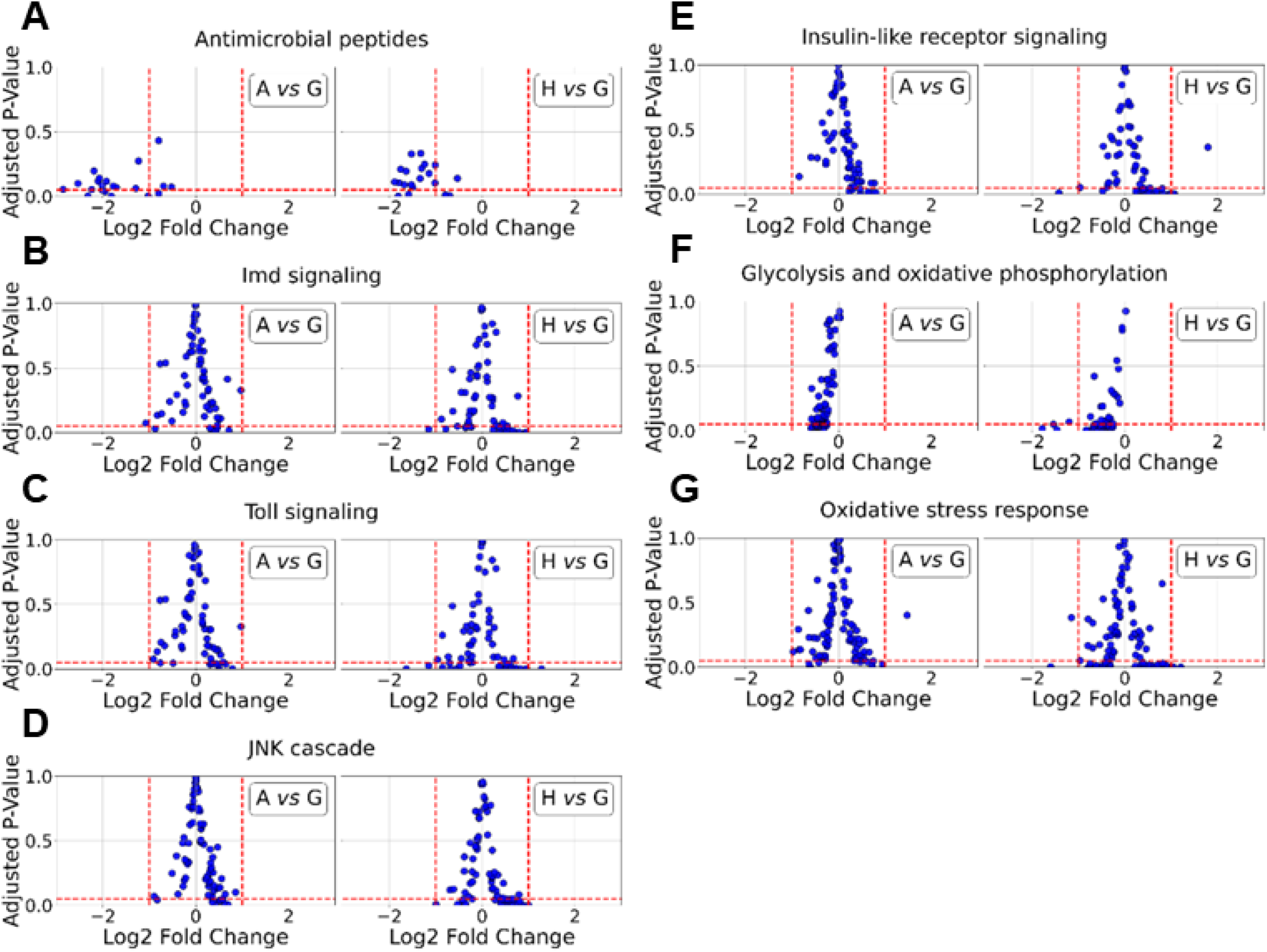
Volcano plots comparing H- and A-to G-conditions for potential cancer-related systemic regulations. (**A-G**) Volcano plots comparing A(left)- and H(right)-to G-conditions for AMPs (**A**), Imd pathway (**B**), Toll pathway (**C**), JNK pathway (**D**), Insulin pathway (**E**), energy metabolism (**F**) and oxidative stress (**G**). Note that the expression of only the AMP-encoding genes (**A**) is significantly reduced in A- and H-*versus* G-flies, while genes related to energy metabolism tend, although not significantly, to be downregulated (**F**).

**Figure 4:**
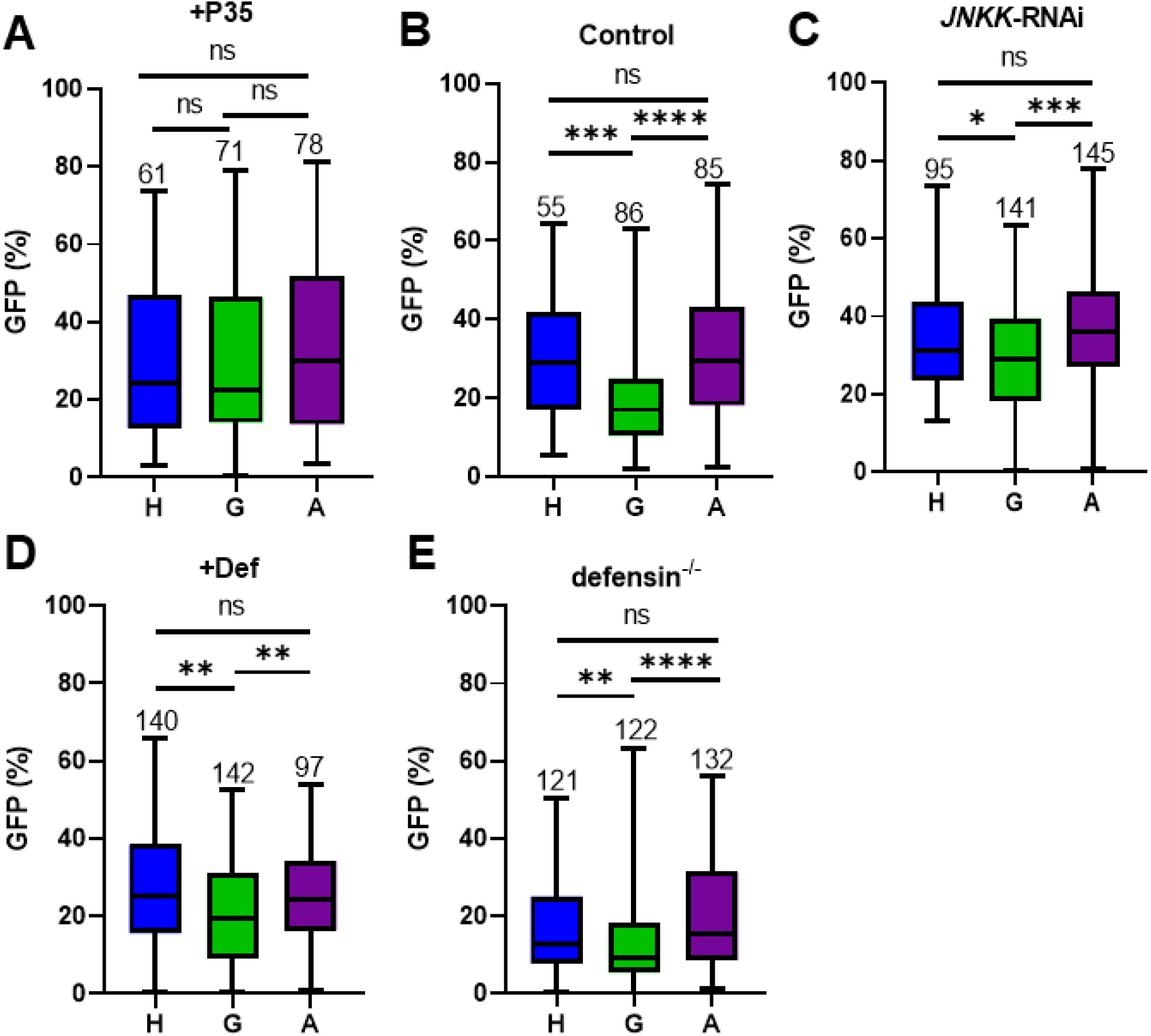
Modulating the social effect on tumor growth. (**A**) Tumor growth in H-, G- and A-tumorous flies either controls (**A**), expressing p35 in tumors (**B**), expressing JNKK-RNAi in tumors (**C**), expressing Defensin in tumors (**D**) or homozygous mutants for *defensin* (**E**). Tumors size is quantified as the percentage of GFP+ cells relative to the total number of intestinal cells. Each sample corresponds to a single gut; sample numbers (n) are indicated above each plot. Note that p35 miss-expression in tumors is the sole genetic setting resulting in the loss of the social-induced tumor suppression.

**Figure 5:**
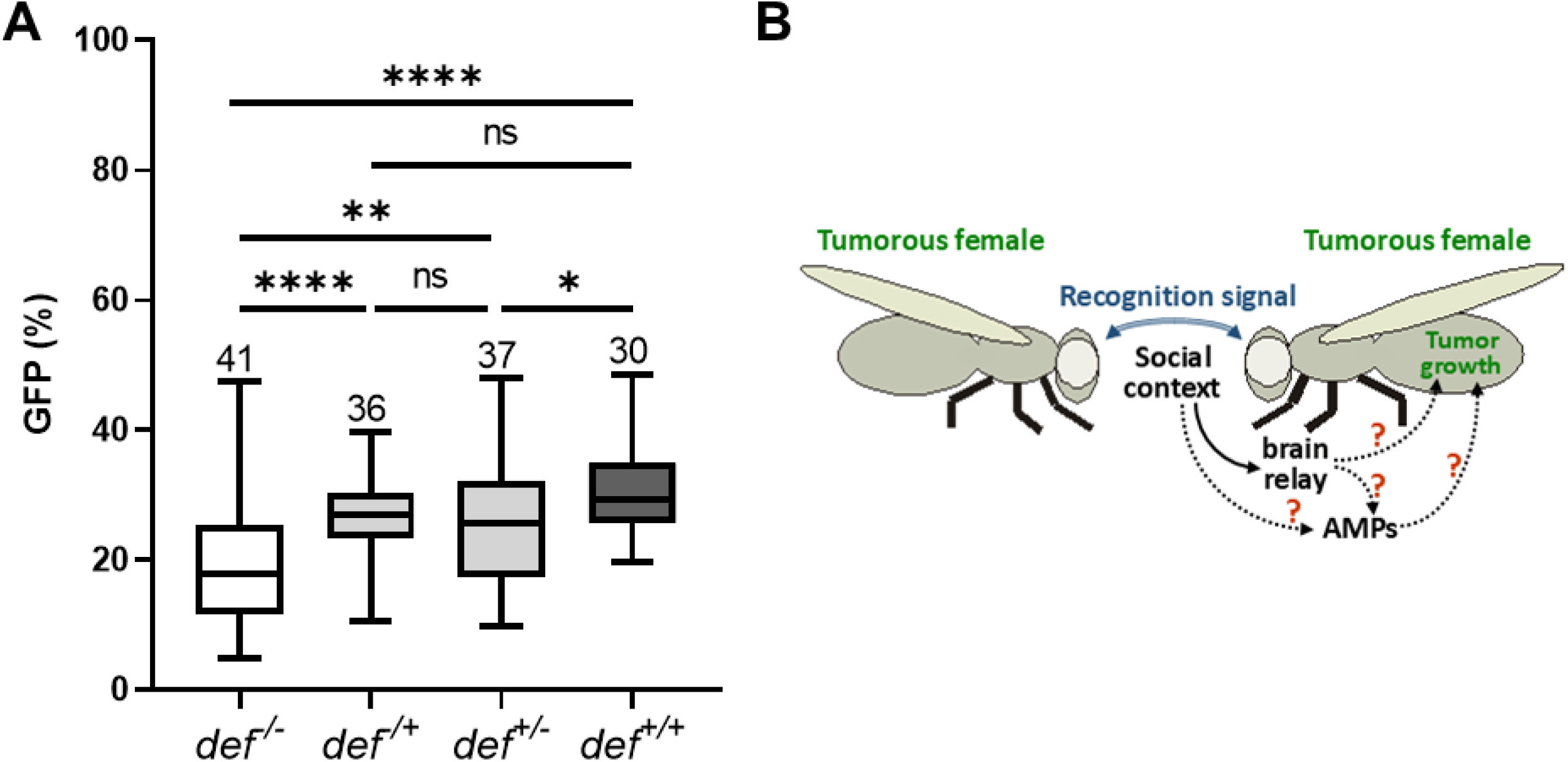
Tumor growth in *defensin* mutant flies. (**A**) The growth of genetically-induced tumors is strongly reduced in homozygous *defensin* mutant flies. A moderate tumor suppression is observed in heterozygous *defensin* mutant flies, which is significant only when the *defensin* mutation is provided by the parental mother flies. Each sample corresponds to a group of two to three guts; sample numbers (n) are indicated under each plot. (**B**) Putative model for tumor growth slowdown in response to social interactions: tumorous flies recognize each other through an as-yet unknown signal, prompting social interactions, which in turn induce changes in the expression of neurological related and AMP-encoded genes. Whether changes in the expression of AMP-encoded genes is directly or indirectly controlled by neurological related genes remains to be investigated.

### The social-induced effect on tumor growth depends on apoptosis but not of JNK signaling and of the immune peptide Defensin

Following the study of Parvy and colleagues showing that the immune peptide Defensin acts through a JNK-signaling to promote apoptosis in imaginal disc tumors (Parvy *et al*. 2019), we investigated whether apoptosis might be responsible of the social-dependent effect. To address this issue, we induced the expression of the baculovirus apoptotic-inhibitor p35 in tumor cells only (Fig. 4A). This apoptosis suppression in tumors resulted in a loss of the social-dependent effect, indicating that cell death is responsible for the lower tumor growth observed in flies maintained in homogenous group (Fig. 4A-B). Next, we analyzed the role of JNK signaling and of the Defensin immune peptide. To address these issues, we expressed in tumor cells only either the Defensin or an interfering RNA (RNAi) to JNKK, which phosphorylates JNK (Glise *et al*. 1995). Interestingly, none of these genetic alterations suppressed the social-dependent effect on tumor growth (Fig. 4C-D). Taken together, these findings indicate that the social-induced tumor suppression depends on apoptosis but neither of the JNK-signaling in tumor cells nor of Defensin. As for most immune peptides, Defensin is mainly synthesized in the fat body to affect immunity and imaginal disc tumor growth in a systemic manner (Lemaitre and Hoffmann 2007; Parvy *et al*. 2018). That Defensin over-expression in tumor cells did not affect social-dependent tumor growth suggests either that Defensin is not responsible for the social-dependent effect or that Defensin was not efficient when produced in tumor cells. Therefore, we generated fly lines to produce tumors in a *defensin* mutant (*def*^*sk3*^) background. In this genetic setting, the social-dependent effect was still observed (Fig. 4E), further validating that the social-induced tumor suppression operates independently of Defensin.

### Defensin sustains the growth of our intestinal tumor model

Surprisingly, the tumor size analyzed in *defensin* homozygote mutants appeared particularly reduced for each social condition (Fig. 4E), suggesting that in the *def*^*sk3*^ mutant background, tumor growth is overall reduced. We therefore, compared the growth of tumors in flies either homozygote, heterozygote or wild-type for the *defensin* gene. Given that our tumorous flies are the progeny of a cross between two parental lines, we had to recombine each parental line with the *def*^*sk3*^ mutation (see materials and methods). We obtained *def*^*sk3*^ mutant tumorous flies when crossing *def*^*sk3*^ mutant parental lines, and control tumorous flies when crossing parental lines wild type for the *defensin* gene. Tumorous flies heterozygous for the *def*^*sk3*^ mutation could be obtained by crossing parental lines, when only one of them is mutant for the *defensin* gene. To prevent some hypothetical maternal effect, we performed reciprocal crosses with either female or male parental *def*^*sk3*^ mutant. Compared to wild type *defensin* tumorous flies, heterozygous *def*^*sk3*^ mutant flies exhibited a moderate suppression of tumor growth, which was significant only in the offspring of female, but not male *def*^*sk3*^ mutant (Fig. 5A). In contrast, in homozygous *def*^*sk3*^ mutants, the tumor growth was strongly reduced as compared to any of the other genetic combinations (Fig. 5A). These findings indicate that in contrast to its antitumoral effect on imaginal disc tumors (Parvy *et al*. 2019), Defensin sustains tumor growth in our intestinal model.

## DISCUSSION

Here, we compared the transcriptome of isogenized *Drosophila* females bearing genetically-induced intestinal tumors raised in different social environments. Despite their genetic identity, we observed that several biological functions are impacted. Given that the genetically-identical tumorous flies grew in indistinguishable conditions until heat-shock induced recombination, the modification in gene expression exclusively relies on the social context in which the flies are maintained during the 10 days past tumor-induction.

In a previous report, we reported a social-dependent effect on the expression of genes related to nervous system activity, suggesting that tumorous flies have a cognitive perception of their social environment (Akiki *et al*. 2024). Consistent with the structure of the central nervous system of adult flies, which comprises a thoracico-abdominal ganglion (Doe 2017), we observed, in the transcriptome of the beheaded flies, changes in the expression of genes specific to the activity of the nervous system. Of major interest is the downregulation of AMP encoding genes directly correlated to the social context, for which, tumor growth is the highest (Dawson *et al*. 2018). In mammals, it has been shown that both the innate and the acquired immune systems can affect cancer progression, directing immunotherapy approaches to fight cancer in humans (Marcus *et al*. 2014; Sharma and Allison 2015b; Iwai *et al*. 2017). To date, we cannot ascertain whether the changes in the expression of genes encoding AMPs ─or alternatively cell-adhesion or energy metabolism─ depend on the activity of the nervous system and in turn directly affects tumor growth (Fig. 5B). Further experiments should be designed to investigate these issues, potentially by manipulating the expression of target genes in the nervous system, while monitoring the expression of putative candidate genes (*e*.*g*. AMPs).

Here, we show that the social-induced tumor suppression is lost when apoptosis is inhibited in tumor cells. These tumors generally develop in the anterior midgut whereas founder clones are widely distributed along the entire midgut (Martorell *et al*. 2014; Dawson *et al*. 2018; Delamotte *et al*. 2025), indicating that numerous clones are likely eliminated after induction. Therefore, the differential social-dependent apoptotic rate might occur all along tumor progression or only at early stages. In contrast to a previous report studying neoplastic imaginal discs (Parvy *et al*. 2019), we observed that overall tumor growth was reduced in *defensin* mutants. This unexpected result raises the question of whether Defensin directly sustains the growth of these intestinal tumors. Alternatively, the organismal physiology might be impacted in loss-of-*defensin* mutants leading to either reducing overall activity of the midgut that is primarily exposed to external injuries or increasing in a feedback loop the expression of other AMPs that could subsequently restrain tumor growth. Future analyzes of the *defensin* mutants will be necessary to address these issues.

In summary, together with our previous reports (Dawson *et al*. 2018; Akiki *et al*. 2024), the present study provides evidence that, although not classified as eusocial insects, fruitflies have a cognitive perception of their social environment, which subsequently influences physiological functions. Importantly, we show that the social context also influences AMP expression levels thereby supporting a potential role of the immune system in controlling tumor growth, an issue that must be extensively investigated in the future.

## Author contributions

JM conceived the study; FM, FMP and JM provided financial support; PD, PA, MP, MdTdT and JM performed experiments; PD, DN, XC and JM analyzed the data; PD and JM wrote the manuscript.

## Data availability

No reagent and no original *Drosophila* strain have been generated for this study. All the fly strains were generated by recombining and/or combining lines from the VDRC stock center or generous gifts from A. Casali, B. Lemaitre and S. Szuplewski. The recombined *Drosophila* stocks will be kept in our collections for a few months, but their availability also requires the agreement of the owners of the initial lines. The data for confirming the conclusions of the article are present within the articles, figures and supplementary data.

## Aknowledgements

We wish to thank A. Casali, B. Lemaitre, S. Szuplewski, VDRC and BDSC for fly stocks and A. Le Rouzic for critical advices in transcriptome analysis. We acknowledge the sequencing and bioinformatics expertise of the I2BC High-throughput sequencing facility, supported by France Génomique (funded by the French National Program “Investissement d’Avenir” ANR-10-INBS-09) and the Imagerie-Gif Core facilities for technical assistance (funded by ANR: ANR-11-EQPX-0029/Morphoscope, ANR-10-INBS-04/FranceBioImaging, ANR-11-IDEX-0003-02/Saclay Plant Sciences).

